## Supplementary file for "Decoding the biogenesis of HIV-induced CPSF6 puncta and their fusion with the nuclear speckle"

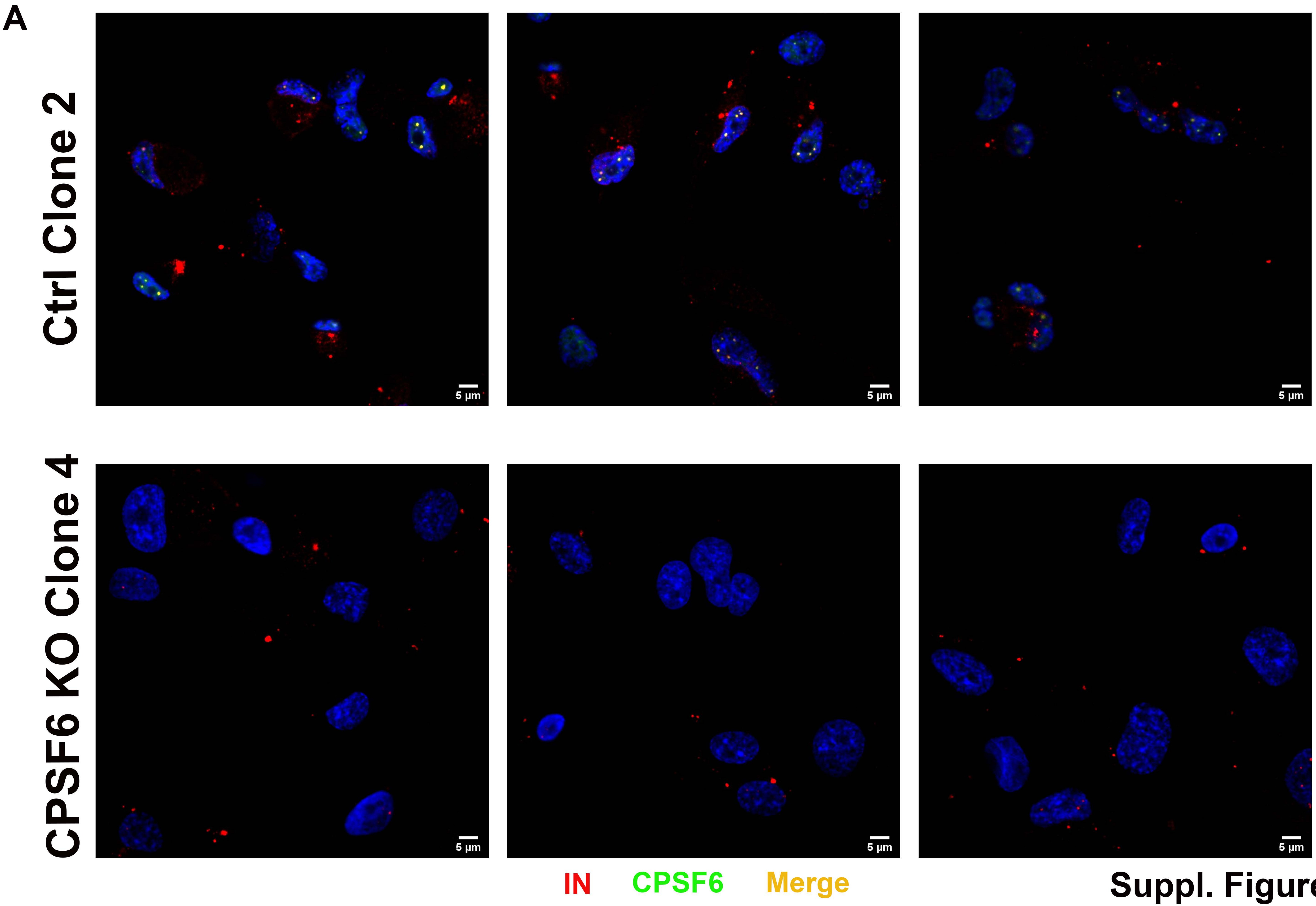


Suppl. Figure 1

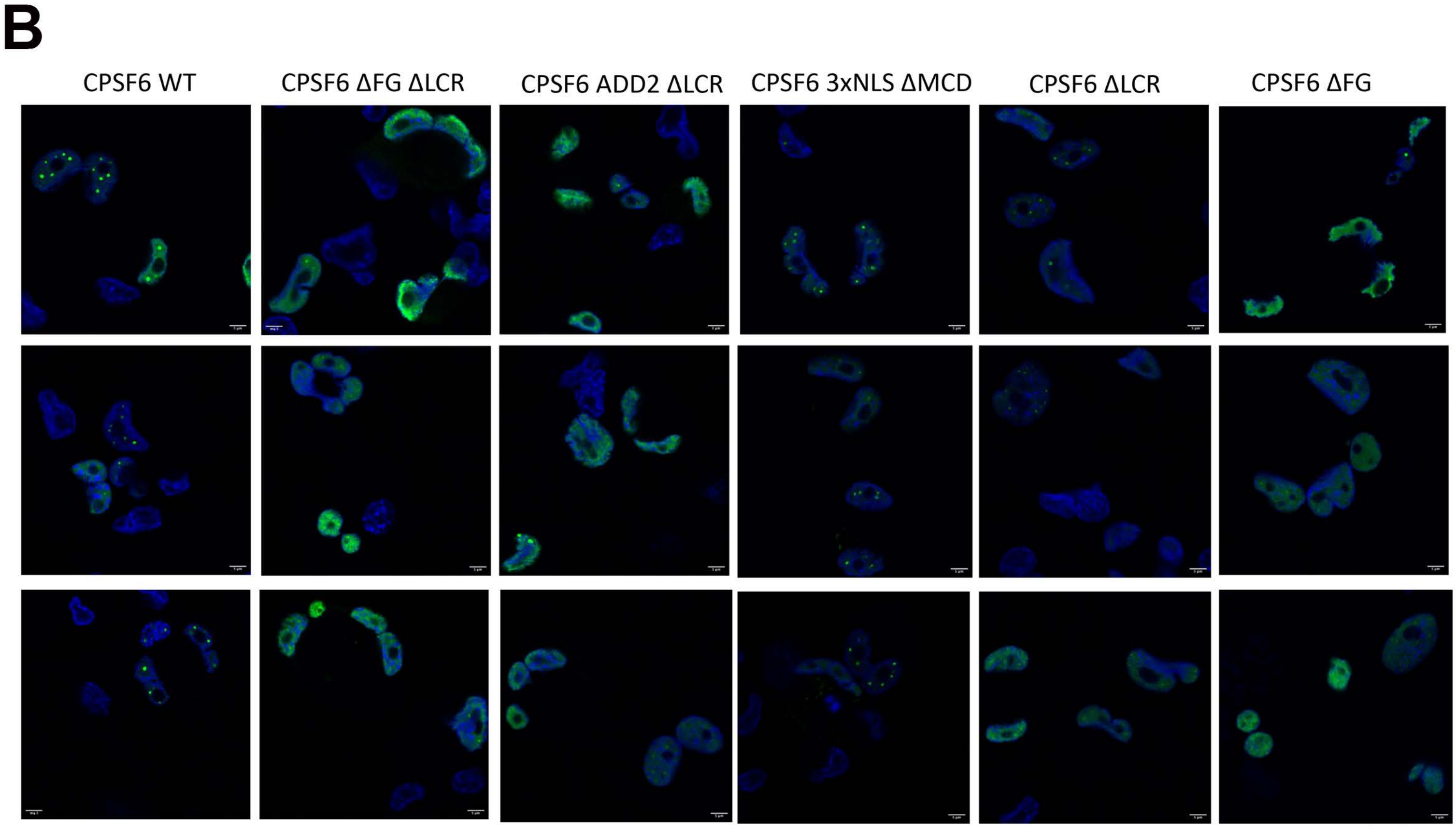


Suppl. Figure 1

A

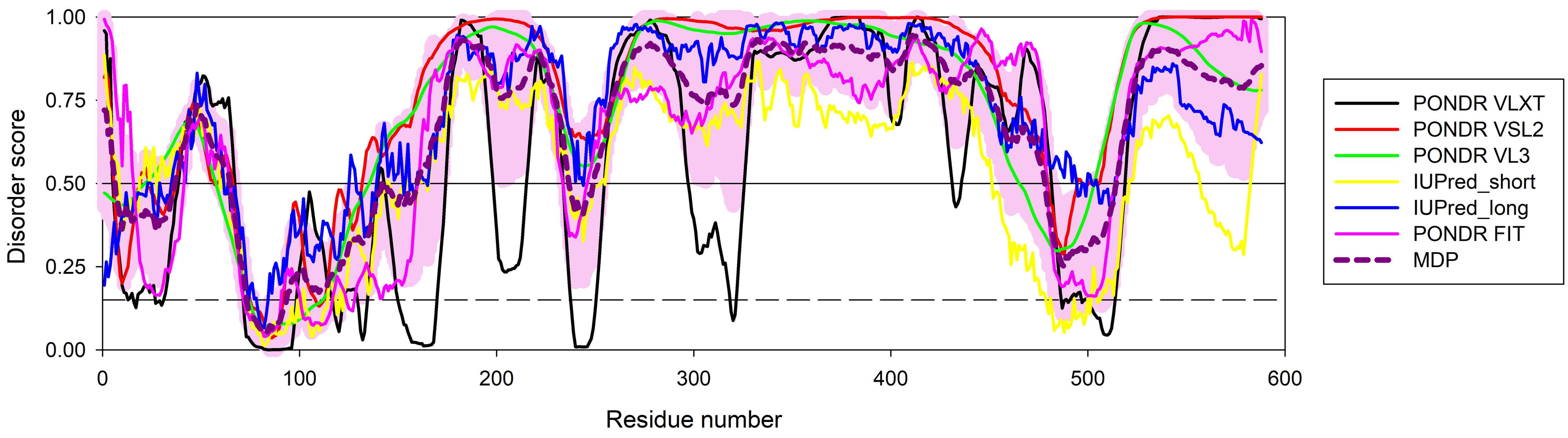


Suppl. Figure 2

B

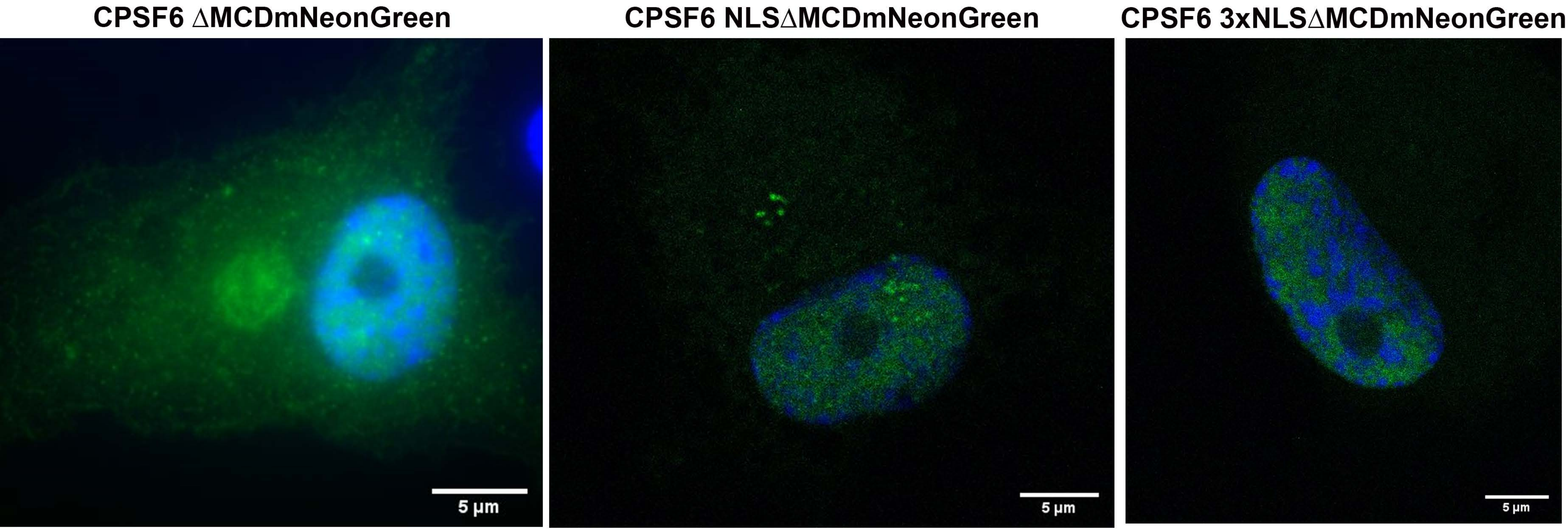


Suppl. Figure 2

**Bioinformatic comparison of amino acids sequence ~261-358aa:**

CPSF6 domain ΔLRC + FG peptide:
gavpggdrfpgpagpggppppfpgnlikhlvkgtrplfletripwhmghsiePVLFPGQPFGQPPLGpfpprppgplgppltlappphlp
gpppgapp

CPSF6 **Δ**LCRs + ADD2 + FG peptide:
DGEKETAPEEPGSPAKSAPASPVQSPAKEAETKSPLVSPSKSLEEGTKKTETPVLFPGQPFGQPPLGSKAATTEPETTQ
PEGVVVNGREEEQTAEEIL

CPSF6 WT (domain LCRs + FG peptide):

Eipifglkagqtpprpplgppgppgppgppppgqvlppplagppnrgdrppppvlfpgqpfgqpplgplppgppppvpgygpppgpppp
qqgpppppg

CPSF6 (domain LCRs **Δ** FG peptide):

Eipifglkagqtpprpplgppgppgppgppppgqvlppplagppnrgdrppppplppgppppvpgygpppgppppqqgpppppg

CPSF6 (**Δ** domain LCRs **Δ** FG peptide):

Pfpprppgplgppltlappphlpgpppgapppaphvnpaffppptnsgmptsdsrgppptdpygrpppydrgdygppgremdtartpls
eaefeeimn

**Suppl. Figure 3**

A

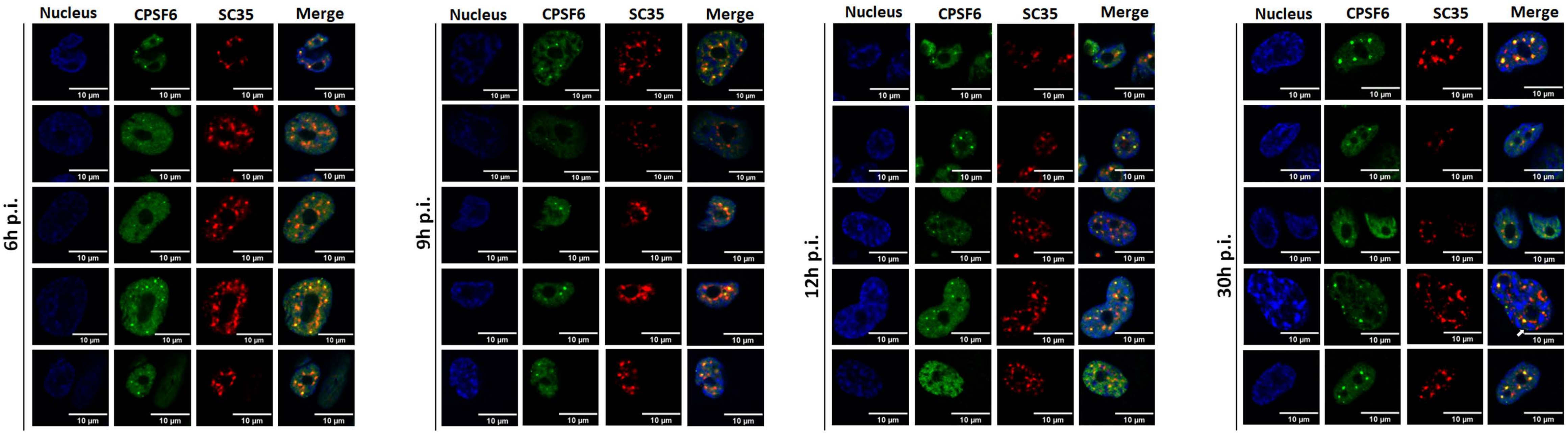


Suppl. Figure 4

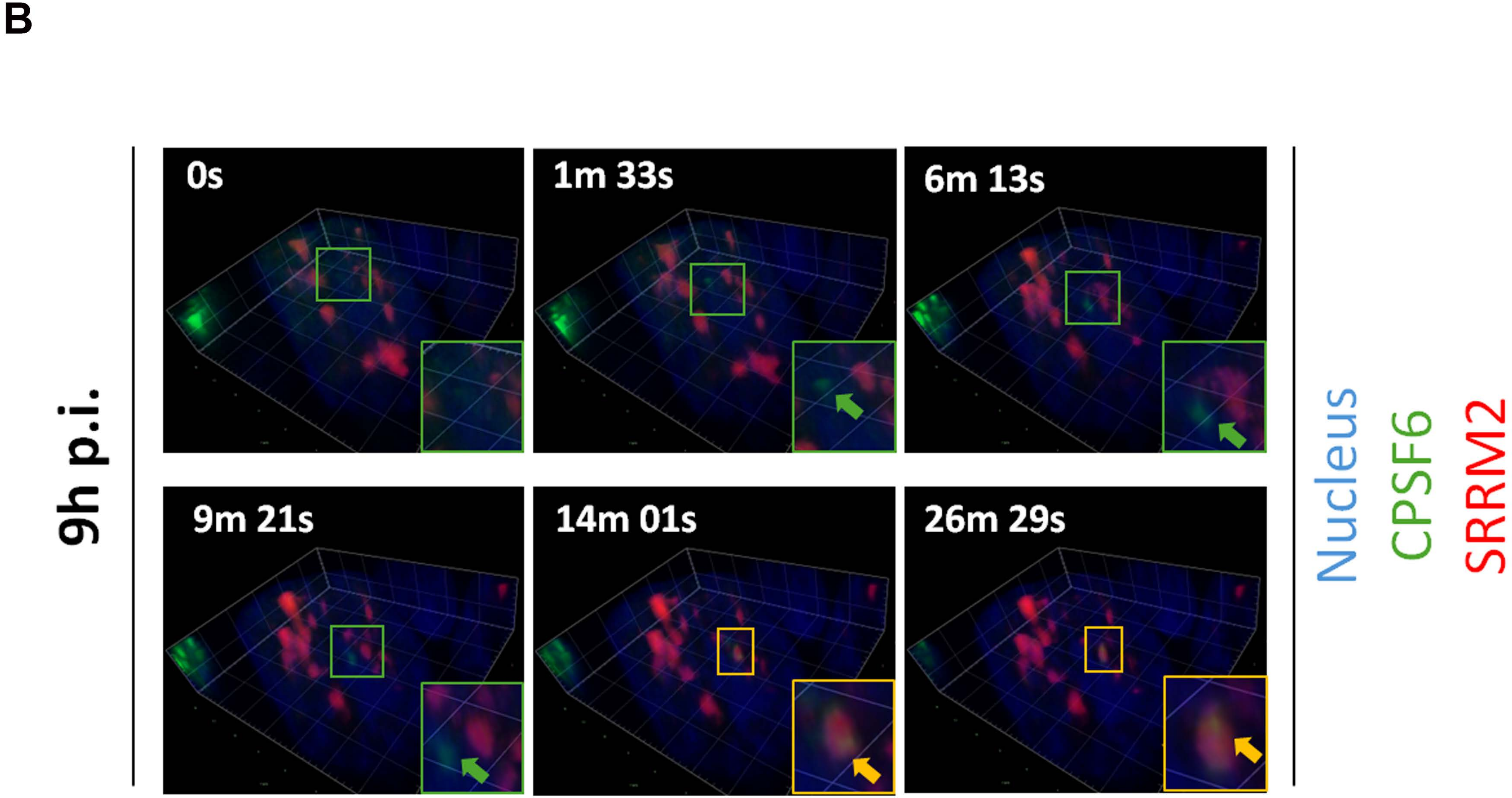


Suppl. Figure 4

A

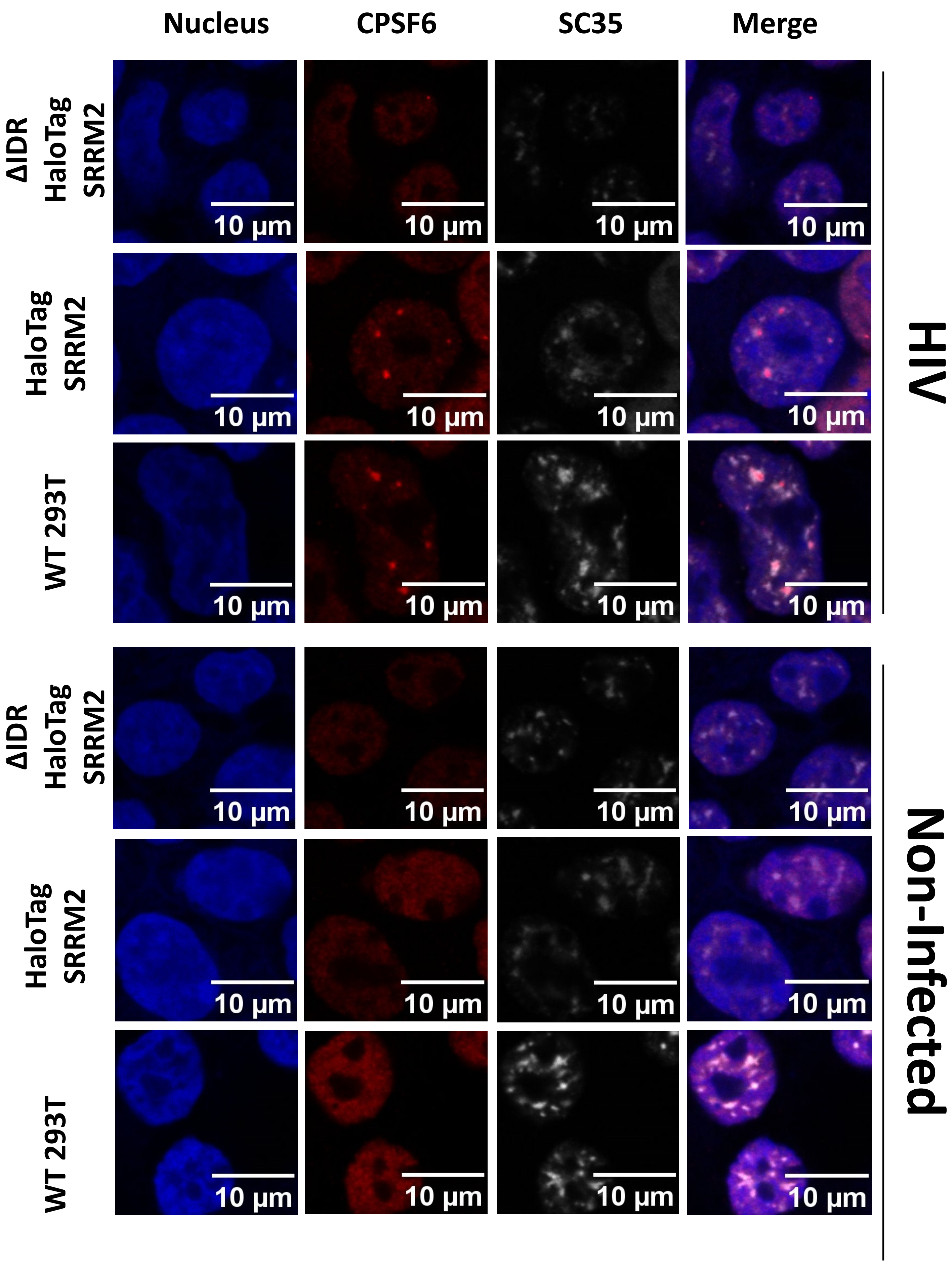

Nucleus
CPSF6
SC35
Merge
ΔIDR HaloTag SRRM2
HaloTag SRRM2
WT 293T
ΔIDR HaloTag SRRM2
HaloTag SRRM2
WT 293T
10 μm
HIV
Non-Infected


Suppl. Fig.5

**B**

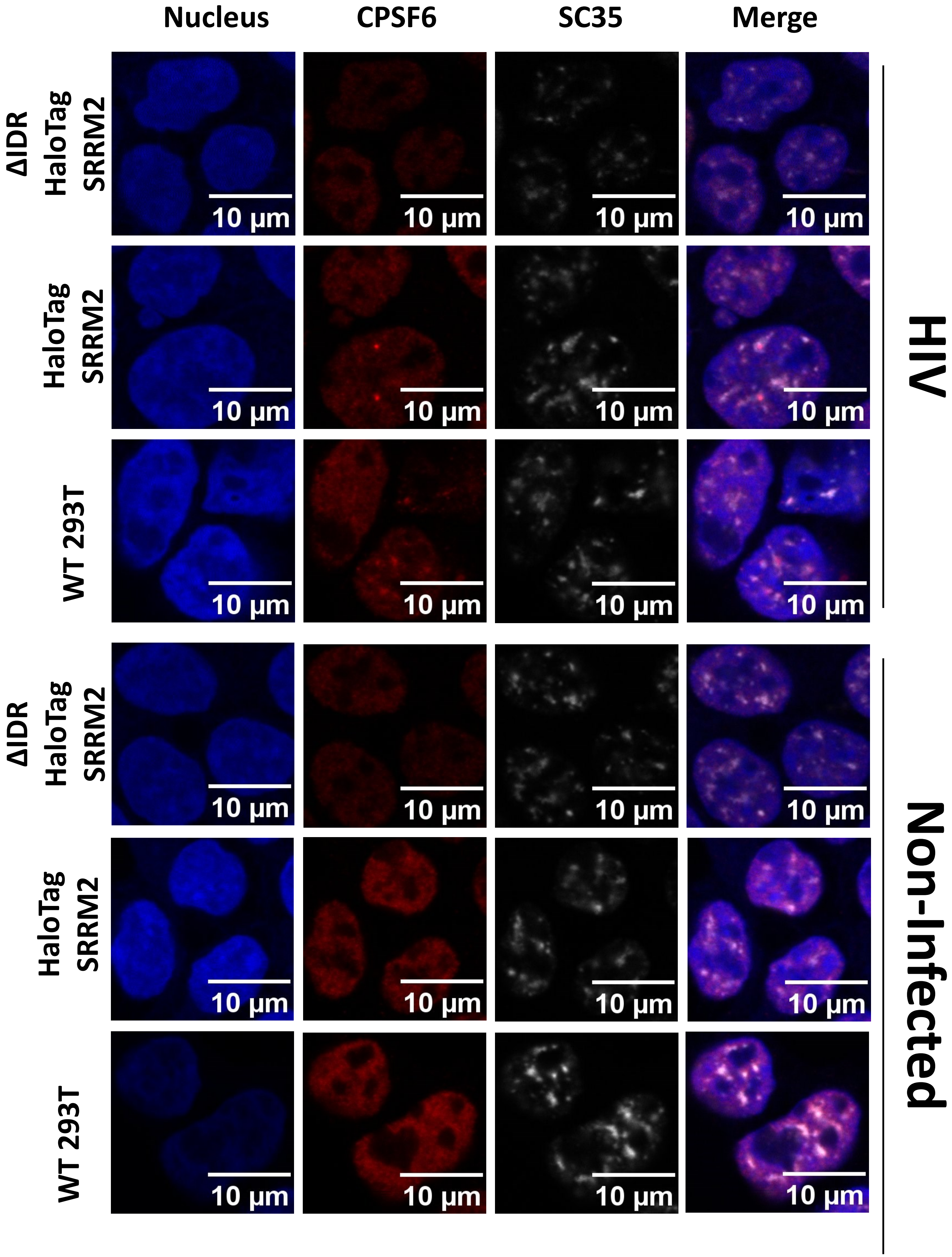
Nucleus
CPSF6
SC35
Merge
ΔIDR HaloTag SRRM2
HaloTag SRRM2
WT 293T
ΔIDR HaloTag SRRM2
HaloTag SRRM2
WT 293T
HIV
Non-Infected
10 µm

Suppl. Fig.5

C

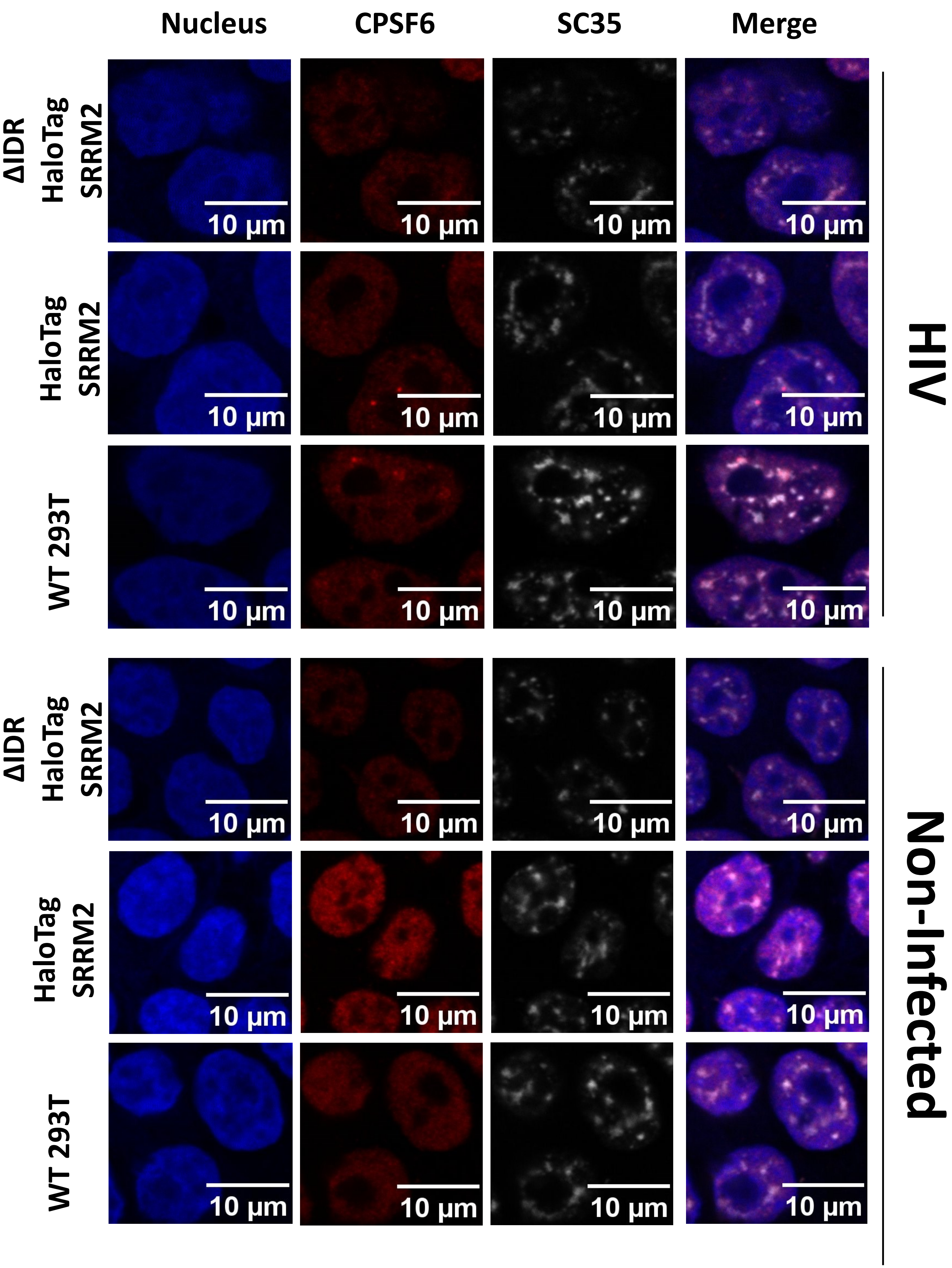
Nucleus
CPSF6
SC35
Merge
ΔIDR HaloTag SRRM2
HaloTag SRRM2
WT 293T
ΔIDR HaloTag SRRM2
HaloTag SRRM2
WT 293T
10 μm
HIV
Non-Infected

Suppl. Fig.5

# D

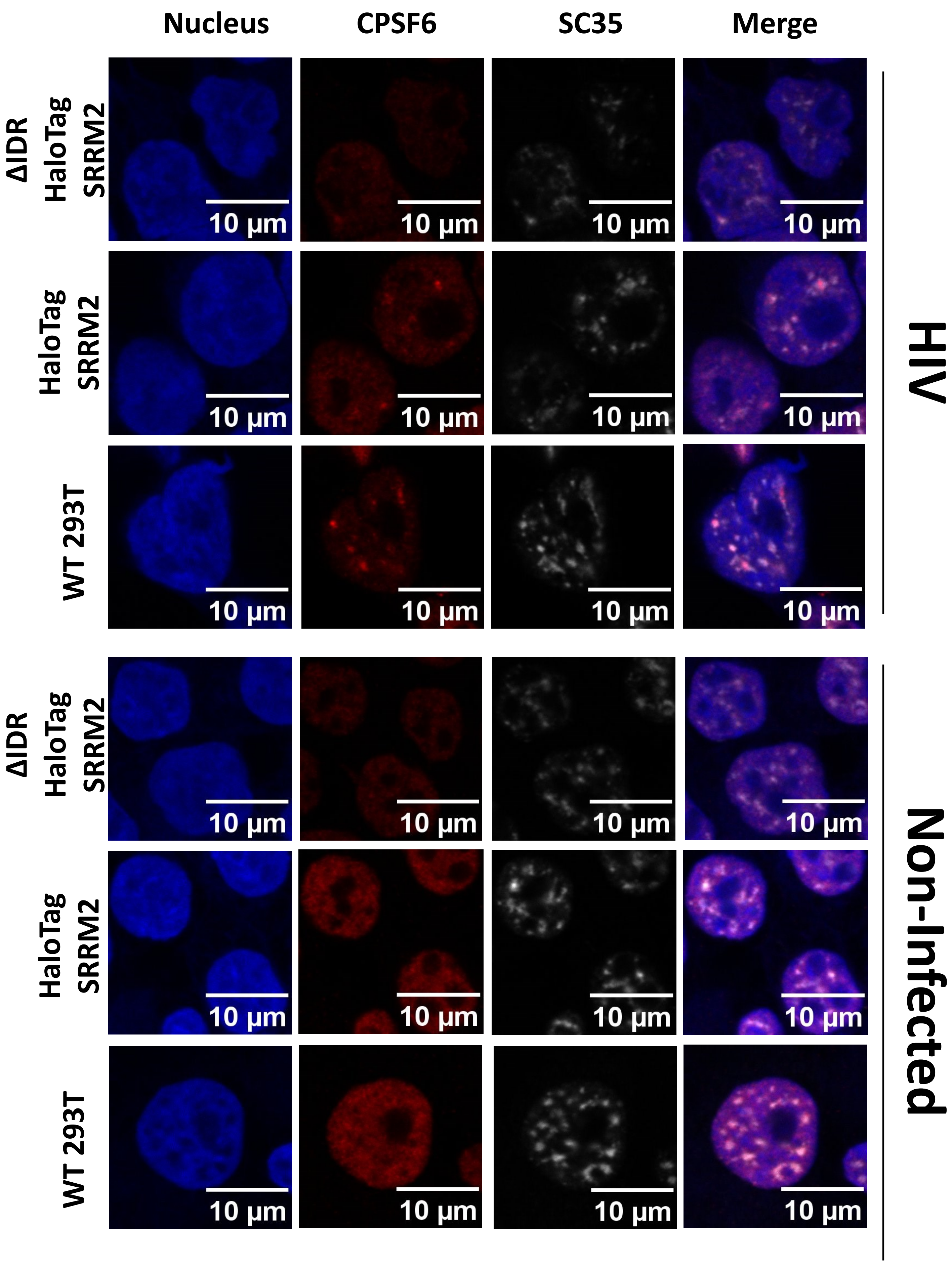

Nucleus
CPSF6
SC35
Merge
ΔIDR HaloTag SRRM2
HaloTag SRRM2
WT 293T
ΔIDR HaloTag SRRM2
HaloTag SRRM2
WT 293T
10 µm
HIV
Non-Infected


Suppl. Fig.5

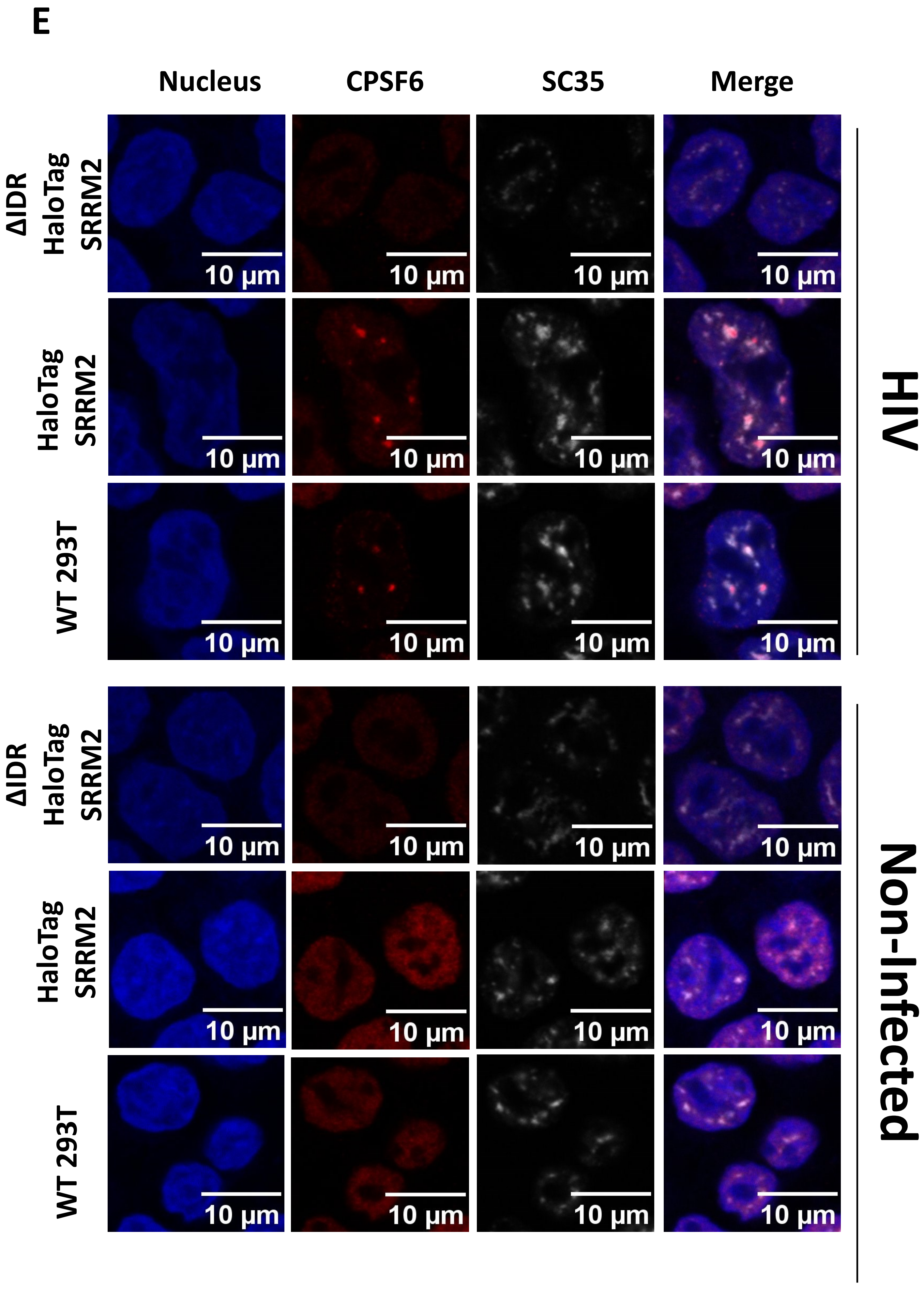
E
Nucleus
CPSF6
SC35
Merge
ΔIDR HaloTag SRRM2
HaloTag SRRM2
WT 293T
ΔIDR HaloTag SRRM2
HaloTag SRRM2
WT 293T
10 μm
HIV
Non-Infected

Suppl. Fig.5

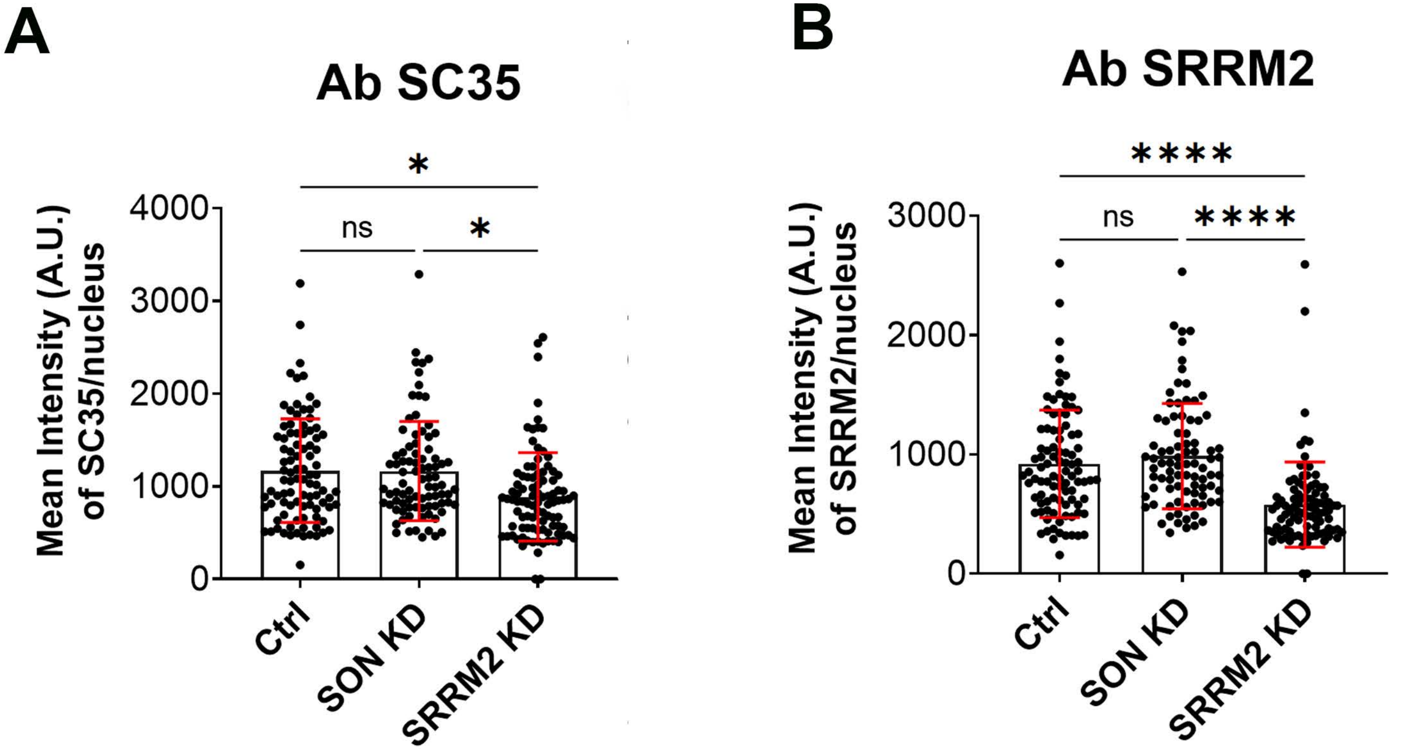
A
Ab SC35
Mean Intensity (A.U.) of SC35/nucleus
4000
3000
2000
1000
0
*
ns
*
Ctrl
SON KD
SRRM2 KD
B
Ab SRRM2
Mean Intensity (A.U.) of SRRM2/nucleus
3000
2000
1000
0
****
ns
****
Ctrl
SON KD
SRRM2 KD

Suppl. Fig.6

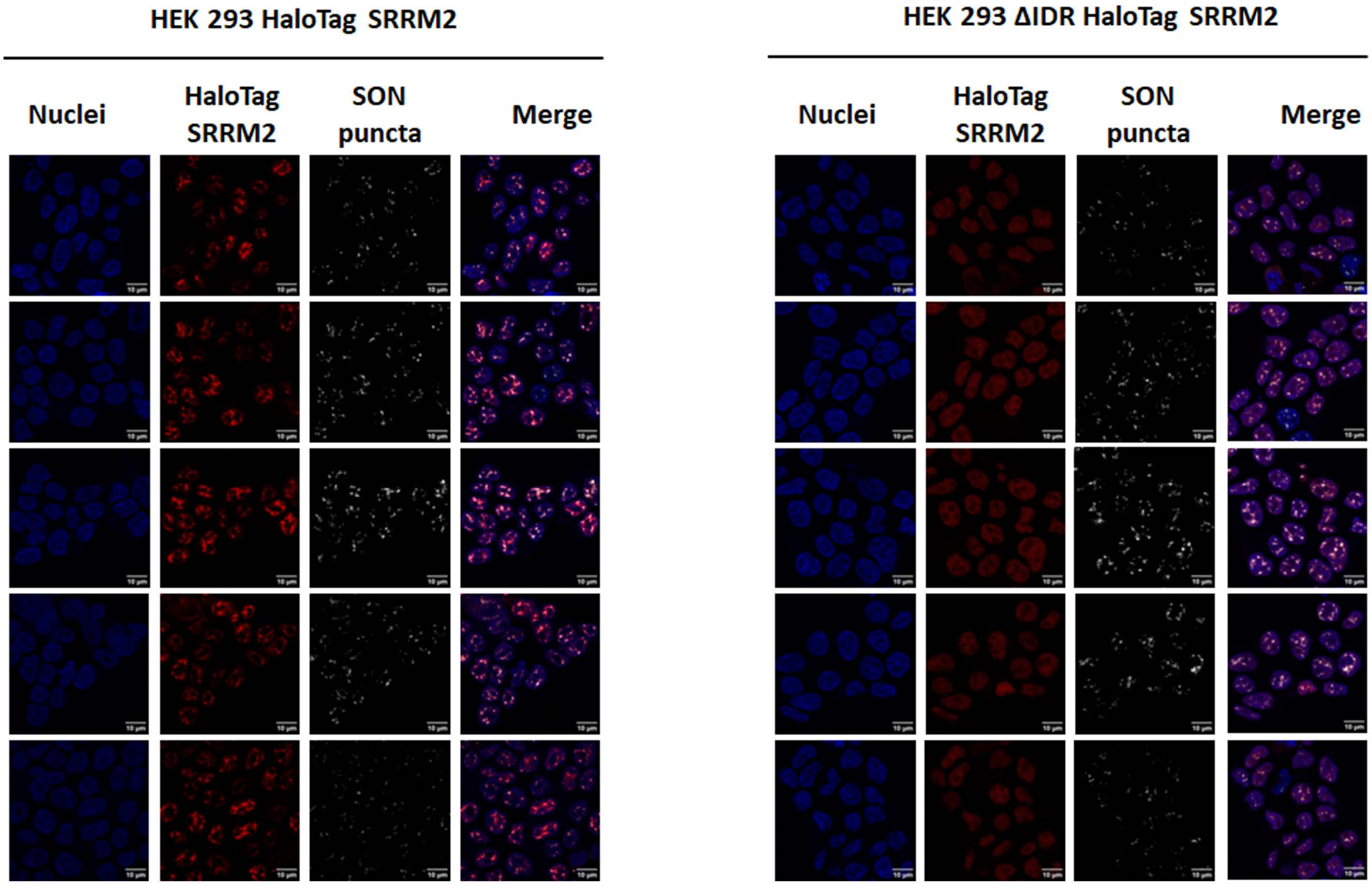


Suppl. Figure 7
